# Serum Amyloid A promotes Acetaminophen-induced liver injury by damaging sinusoidal endothelial cell and exacerbating platelet aggregation in liver

**DOI:** 10.1101/2021.12.01.470869

**Authors:** Kai You, Yan Wang, Xiaoxia Chen, Zhen Yang, Yan Chen, Shenglin Tan, Jiawang Tao, Anteneh Getachew, Tingcai Pan, Yingying Xu, Yuanqi Zhuang, Fan Yang, Xianhua Lin, Yinxiong Li

## Abstract

**Background:** Acetaminophen (APAP) is the most commonly used non-prescription antipyretic and analgesic drugs. Overuse of APAP can cause hepatotoxicity. Liver sinusoidal endothelial cells (LSECs) damage is an important early event in APAP-induced liver injury. Serum amyloid A (SAA) is an acute phase protein that mainly produced by hepatocytes, and promotes endothelial dysfunction via a pro-inflammatory and pro-thrombotic effect in atherosclerosis and renal disease. However, the role of SAA in APAP-induced liver injury remains unclear.

**Methods:** In this study, we used neutralizing antibody (anti-SAA) or antagonistic small peptide derived from sequence of human SAA1/2 (SAA-pep) to block the functional activity of Saa1/2 in mouse serum. Immunohistochemistry staining, Evans blue and platelet adhesion assays were performed to examine the liver damage, the integrity of sinusoidal endothelium and platelets accumulation in APAP-induce liver injury.

**Results:** Our study showed that in the early stage of APAP-induced acute liver injury in mice, the intrahepatic and serum Saa1/2 levels were significantly increased within 24 hours, and then gradually reduced to normal level from 3 days. Neutralization of Saa1/2 by antibodies or peptides effectively prevented the destruction of hepatic sinusoids, reduced the intrahepatic hemorrhage and platelet accumulation in liver, as well as increased the survival rate of mice treated with lethal dose of APAP. *In vitro* experiments showed that Saa1/2 aggravated LSECs death induced by APAP. Moreover, Saa1/2 promoted platelets adhesion on LSECs via Tlr2/Vcam-1 axis.

**Conclusion:** Our findings suggest that Saa1/2 promotes APAP-induced liver injury by damaged LSECs and exacerbated platelets aggregation. This study provides a potential target for intervention of acute liver injury/failure caused by hepatotoxic drugs such as APAP.

## Introduction

Acetaminophen (APAP) is the most commonly used over-the-counter drug for the treatment of pain and fever. Meanwhile, APAP is also a dose-related toxin which can lead to liver injury/failure in all mammalian species. APAP toxicity dwarfs all other prescription drugs as a cause for acute liver failure (ALF) in United State and Europe, responsible for 46% of all ALF and nearly 500 deaths annually in US, and between 40 and 70% of all ALF cases in Britain and Europe [1].

Normally, APAP is eliminated by glucuronidation and sulfation catalyzed by UDP-glucuronosyl transferases (UGT) and sulfotransferase (SULT) [2]. Hepatotoxicity occurs when overdose of APAP are used. Excess APAP undergoes oxidation to form the highly reactive toxic metabolite N-acetyl-para-benzoquinone imine (NAPQI) by hepatic cytochrome CYP2E1 (to a lesser extent with CYP1A2 and 3A4). Excessive NAPQI can deplete cellular glutathione and bind to a number of cellular proteins, especially mitochondrial proteins, which leads to mitochondrial dysfunction and hepatocyte necrosis [2-4]. Importantly, Liver sinusoidal endothelial cells (LSECs) injury is an important early event in the development of APAP-induced liver injury. LSECs injury is priors to hepatocellular injury as early as 30 minutes after the administration of APAP [5-7]. However, the mechanisms of such hepatic sinusoids injury induced by APAP remain to be uncovered.

Serum amyloid A (SAA) is an acute phase protein that bound to high density lipoproteins (HDL) in the blood. SAA has several variants which are secreted mainly by hepatocytes [8]. SAA1 and SAA2 are inducible by inflammatory cytokines, and its concentration can be increased up to 1000-fold during inflammation, and persistently high levels are evident in chronic pathologies such as diabetes mellitus, rheumatic disorders, cancers and atherosclerosis [9]. SAA3 is thought to be a pseudogene in human, while SAA4 is constitutively produced [8, 10, 11]. Although SAA is mainly produced in the liver, however, the effect of SAA in the pathogenesis of acute liver injury is still unknown. In addition, conclusions from two different studies are conflicting. For instance, one study had showed that SAA displayed a protective role by reduced multifocal necrosis in the liver when pretreatment of SAA prior to Concanavalin A (Con A) administration [12]. However, another study found that overexpression of SAA in mice promoted hepatocyte necrosis and increased CD4 T cells and macrophages infiltration and secretion of inflammatory factors in Con A-induced liver injury [13]. Therefore, the mechanism of SAA on regulation of acute liver injury needs to be investigated.

Recently, it has been found that serum acute phase reactants, including SAA, were significantly upregulated in healthy individuals who are susceptible to APAP-induced liver injury [14]. Serum SAA could also serve as a sensitive biomarker for early detection of hepatotoxicity induced by Ritodrine (a tocolytic agent that acts on β2 adrenoceptor) [15]. The results suggested that SAA might participate in the pathogenesis of drug-induced liver injury. Moreover, it has also been reported that SAA can impair endothelial function by eliciting a pro-inflammatory and pro-thrombotic phenotype that initiate the development of atherosclerosis and renal dysfunction [9, 16]. Based on these findings, we speculated that SAA may potentially involve in regulating early hepatic microvascular dysfunction in APAP-induced liver injury.

## Results

### Saa1/2 was upregulated in the early stage of APAP-induced acute liver injury

In order to study the role of Saa1/2 in APAP-induced liver injury, we firstly determined the expression levels of Saa1/2 in liver tissue and blood after APAP injection. Our data showed that the mRNA and protein levels of Saa1/2 in liver tissues were significantly increased and reached peak at 24 hours after APAP injection (Figure 1A to C). Similarly, ELISA assay showed that the serum level of Saa1/2 was significantly upregulated within 24 hours after injury and gradually decreased to normal level within 7 days (Figure 1D). In addition, to investigate whether APAP could induce Saa1/2 expression in hepatocytes, we isolated primary hepatocytes from normal mice and treated with APAP *in vitro*. We found that treatment of APAP for 3 hours, but not 6 hours, significantly induced the expression and secretion of Saa1/2 protein in hepatocytes (Figure 1E and F). Moreover, the expression of Saa1/2 in hepatocytes was induced by recombinant Saa1/2 proteins in a dose and time dependent manner (Figure 1G). However, the combined treatment of APAP and recombinant Saa1/2 for 6 hours did not induce more Saa1/2 expression than that treated with recombinant Saa1/2 alone. These findings suggest that APAP may induce the expression and secretion of Saa1/2 in hepatocytes at the early stage of APAP-induced acute liver injury.

**Figure 1.**
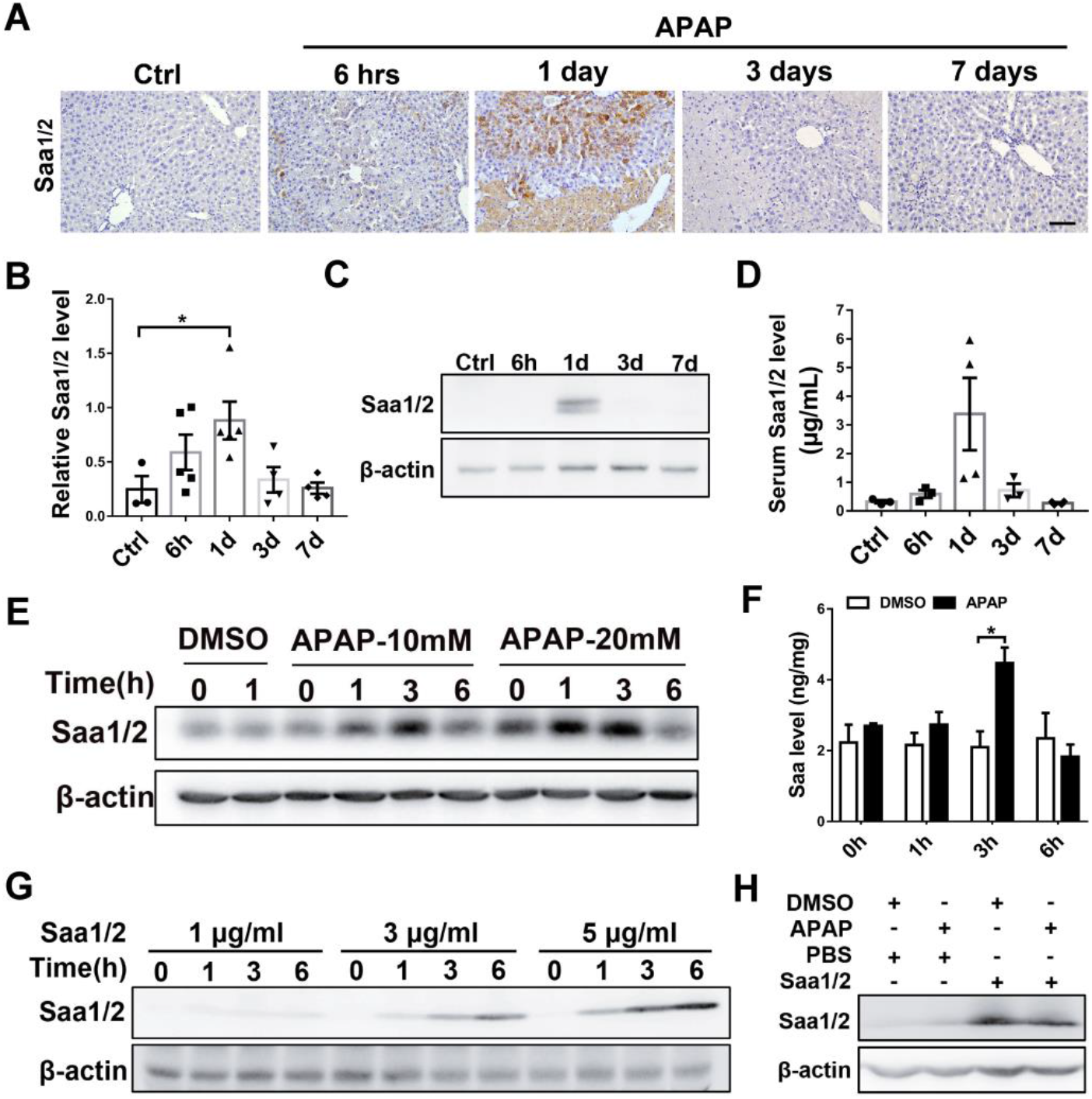
The expression of Saa1/2 was upregulated in the early stage of APAP-induced liver injury. Mice were intraperitoneally injected with APAP (300 mg/kg), the liver tissues and serum were collected for further analysis. (A) Immunohistochemistry staining of Saa1/2 in liver sections; (B-C) Expression of Saa1/2 in liver tissues were detected by q-PCR (n=3-5) (B) or western blots (C); (D) The Saa1/2 level in serum was detected by ELISA (n=3-4); (E-F) Primary hepatocytes were isolated from normal mice that were treated with APAP (10 or 20 mM) for indicated times, the intracellular Saa1/2 level was detected by western blots (E), and the secretive Saa1/2 in hepatocyte culture medium was detected by ELISA (F) (n=3); (G) Primary hepatocytes were treated with Saa1/2 (mixture of Saa1 and Saa2) recombinant proteins for the indicated times, and Saa1/2 expression was measured by western blots. (H) Primary hepatocytes were treated with APAP (10 mM) or Saa1/2 (3 μg/ml) alone or mix together for 6 hours, and Saa1/2 level was measured by western blots (H). Scale bar = 75 μm. *, *p < 0*.*05*.

### Inhibition of Saa1/2 reduced APAP-induced liver injury

In order to further study the functional effect of Saa1/2 on APAP-induced liver injury, we firstly used neutralizing antibody to block the Saa1/2 function before APAP injection (Figure 2A), and then to determine the degree of liver injury and the survival rate of mice. Our data showed that neutralizing Saa1/2 by anti-SAA antibody significantly reduced the activities of ALT and AST in serum (Figure 2B) and the necrotic area in liver (Figure 2C and D). Moreover, the survival rate of mice treated with lethal dose of APAP (500 mg/kg) in the anti-SAA group were significantly higher than that in the IgG (control) group (Figure 2E, n=15). In addition, it has been reported that a small peptide homologous to human SAA1 protein (aa29-42) could inhibit the function of both human and mouse Saa1/2. Therefore, we synthesized this small peptide (named SAA-pep) and the Scramble peptide to study their effects on APAP-induced liver injury (Figure 2F). Our results showed that SAA-pep reduced the activity of ALT in serum at 6 hours after APAP injection (Figure 2G), and significantly decreased the necrotic area of injured liver induced by APAP (Figure 2H and I). Altogether, these findings suggest that inhibition of Saa1/2 may effectively attenuate liver injury and improve the survival of mice injured by APAP injection.

**Figure 2.**
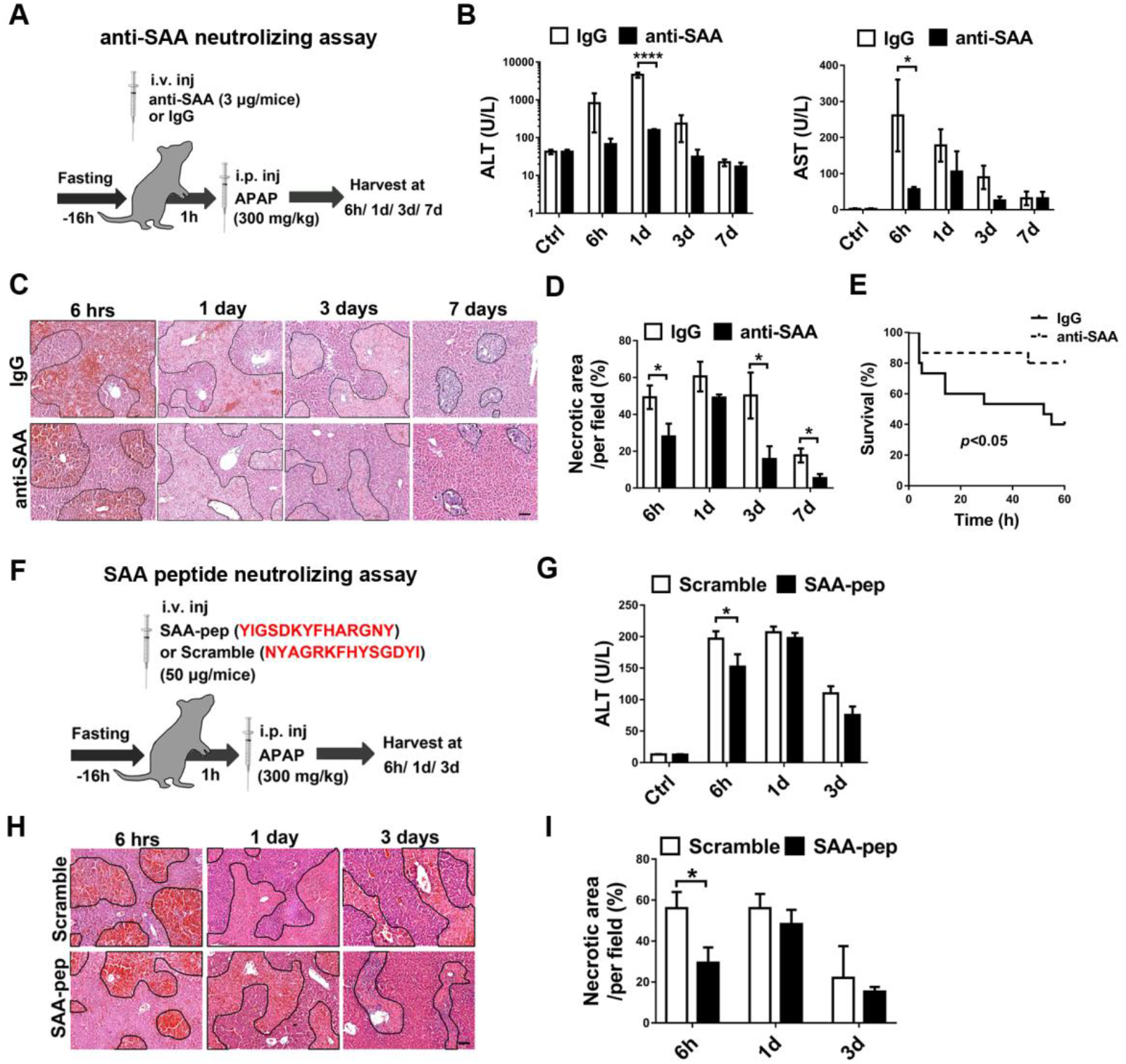
Neutralizing Saa1/2 alleviated APAP-induced liver injury. (A) Schematic of mice receiving anti-SAA neutralizing antibodies treatment one hour before APAP-induced mouse liver injury; (B) serum concentrations of ALT and AST from mice treated with IgG or anti-SAA neutralizing antibodies (n=3-4); (C) H&E staining images (scale bars = 100 μm); (D) quantification of liver necrosis areas in H&E staining sections (n=3-7); (E) Survival curves of mice treated with either IgG or neutralizing antibodies of Saa1/2 (anti-SAA, 5 μg/mice) one hour before intraperitoneal injection of APAP (500 mg/kg) (n = 15); (F) Schematic of mice receiving neutralizing peptide derived from human SAA1 (aa29-42) or the scramble peptide (50 μg/mice) one hour before APAP-induced mouse liver injury; (G) serum samples were collected for detection of ALT activities (n=3-4); (H) the liver tissues were collected for H&E staining (Scale bar = 100 μm), (I) quantification of necrosis areas in H&E staining sections (n=3-7). *, *p* < 0.05; ****, *p* < 0.001.

### Inhibition of Saa1/2 reduced intrahepatic hemorrhage and platelet aggregation

It has been known that hepatic microvascular injury was an early event in the pathogenesis of APAP-induced liver injury, as shown by intrahepatic hemorrhage and congestion precede the appearance of hepatocyte necrosis. According to our results from gross anatomy of livers and H&E staining, we found that neutralizing Saa1/2 significantly alleviated APAP-induced intrahepatic hemorrhage (Figure 3A and B). Furthermore, immunofluorescence staining of SE-1, which is a specific hepatic sinusoidal endothelial cell antibody, was performed in liver section. We found that neutralization of Saa1/2 maintained the integrity of hepatic microvascular structure as compared with IgG control mice after 6 hours of APAP treatment (Figure 3C). To further investigate the effect of Saa1/2 on the permeability of sinusoidal endothelium, we intravenously injected Evans Blue dye into mice that were treated with or without anti-SAA in APAP model, and found that neutralizing Saa1/2 significantly reduced the accumulation of Evans Blue dye in the liver (Figure 3D and E). These data suggest that Saa1/2 may promote the hepatic microvascular injury in the early stage of APAP-induced liver injury. In addition, we further to determine the effect of Saa1/2 on the intrahepatic accumulation of platelets after APAP treatment. Our data showed that blockage of the Saa1/2 function by neutralizing antibody or SAA-pep both significantly decreased intrahepatic platelet aggregation at the injury site at 6 hours after APAP treatment (Figure 3F and G, Supplementary Figure S1). Altogether, these findings suggest that Saa1/2 may promote hepatic sinusoidal endothelial cells injury, platelets accumulation in liver and increase the permeability of sinusoidal endothelium, which results in aggravation of liver injury induced by APAP.

**Figure 3.**
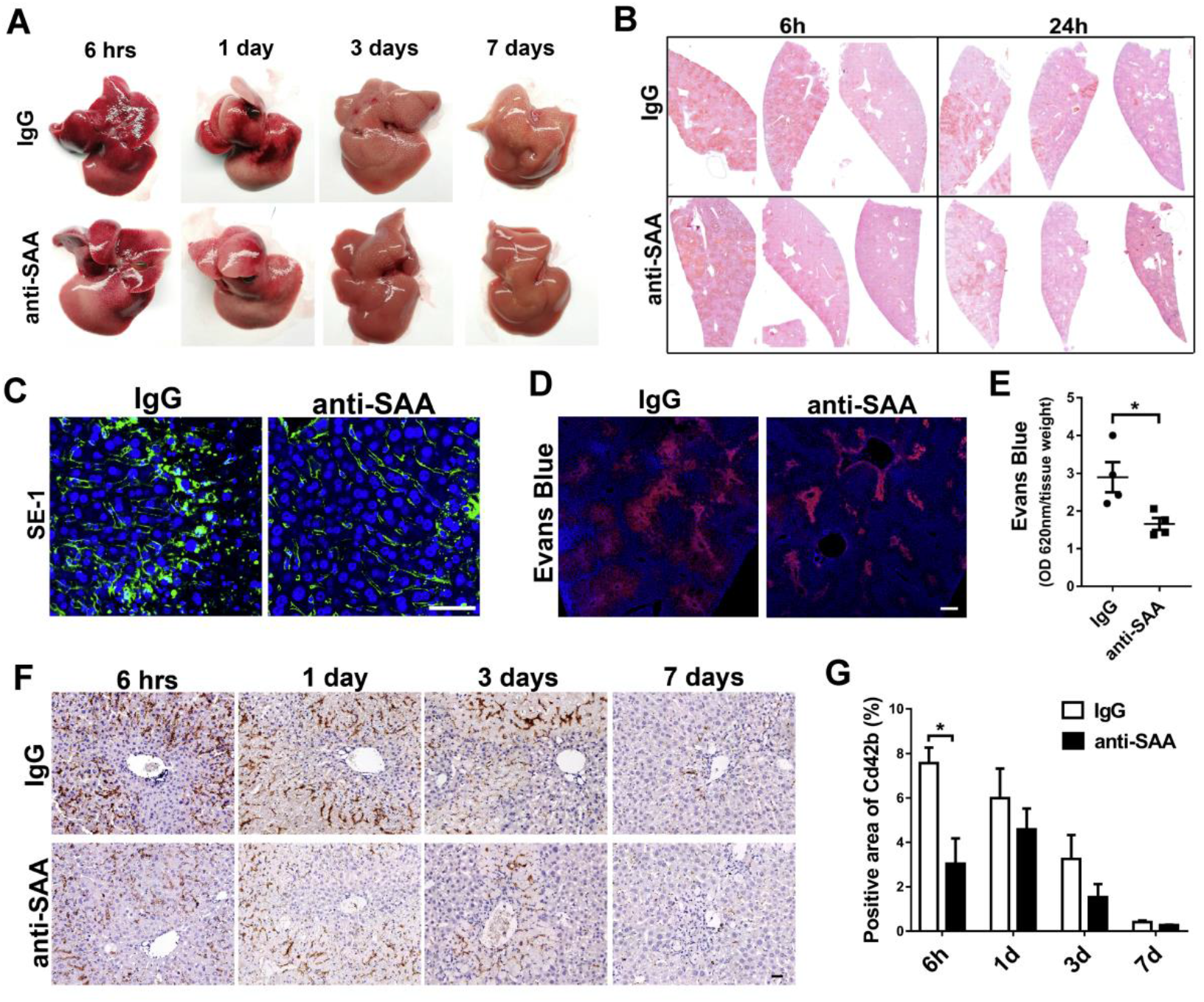
Inhibition of Saa1/2 maintained the integrity of hepatic sinusoids and reduced intrahepatic hemorrhage and platelet aggregation. Mice were treated with IgG or anti-SAA (3 μg/mice) one hour before intraperitoneal injection of APAP (300 mg/kg). (A) Representative gross anatomy of livers; (B) Representative H&E staining images of liver; (C) Immunofluorescence staining of liver sinusoidal endothelial cell with SE-1 antibody (scale bars = 50 μm); (D) Representative images of Evans Blue assay (scale bars = 200 μm), (E) and the concentration of Evans Blue dye in liver tissue homogenate were detected by spectrophotometer at 620 nm (n=4); (F) Immunohistochemistry staining of platelet with Cd42b antibody (scale bars = 50 μm), (G) and quantitative data of Cd42b positive areas in (F) (n=3-4). *, *p* < 0.05.

### Saa1/2 enhanced the injury effect of APAP on LSECs

In order to explore the mechanism of Saa1/2 in promoting intrahepatic hemorrhage, primary LSECs were isolated from mice for study (Supplementary Figure S2). Firstly, flow cytometry analysis was performed to determine the proportion of dead LSECs treated with APAP, we found that the death of LSECs was significantly increased by APAP in a dose dependent manner (Figure 4A and B). Subsequently, LSECs were treated by recombinant Saa1/2 combined with a lower concentration of APAP (10 mM), which is equivalent to the *in vivo* concentration of APAP after injection at a dose of 300 mg/kg. Flow cytometry analysis and Hoechst 33342/PI double staining results showed that Saa1/2 and APAP synergistically induced LSECs death (Figure 4C to F). These findings suggest that Saa1/2 may enhance the susceptibility of LSECs to APAP toxicity.

**Figure 4.**
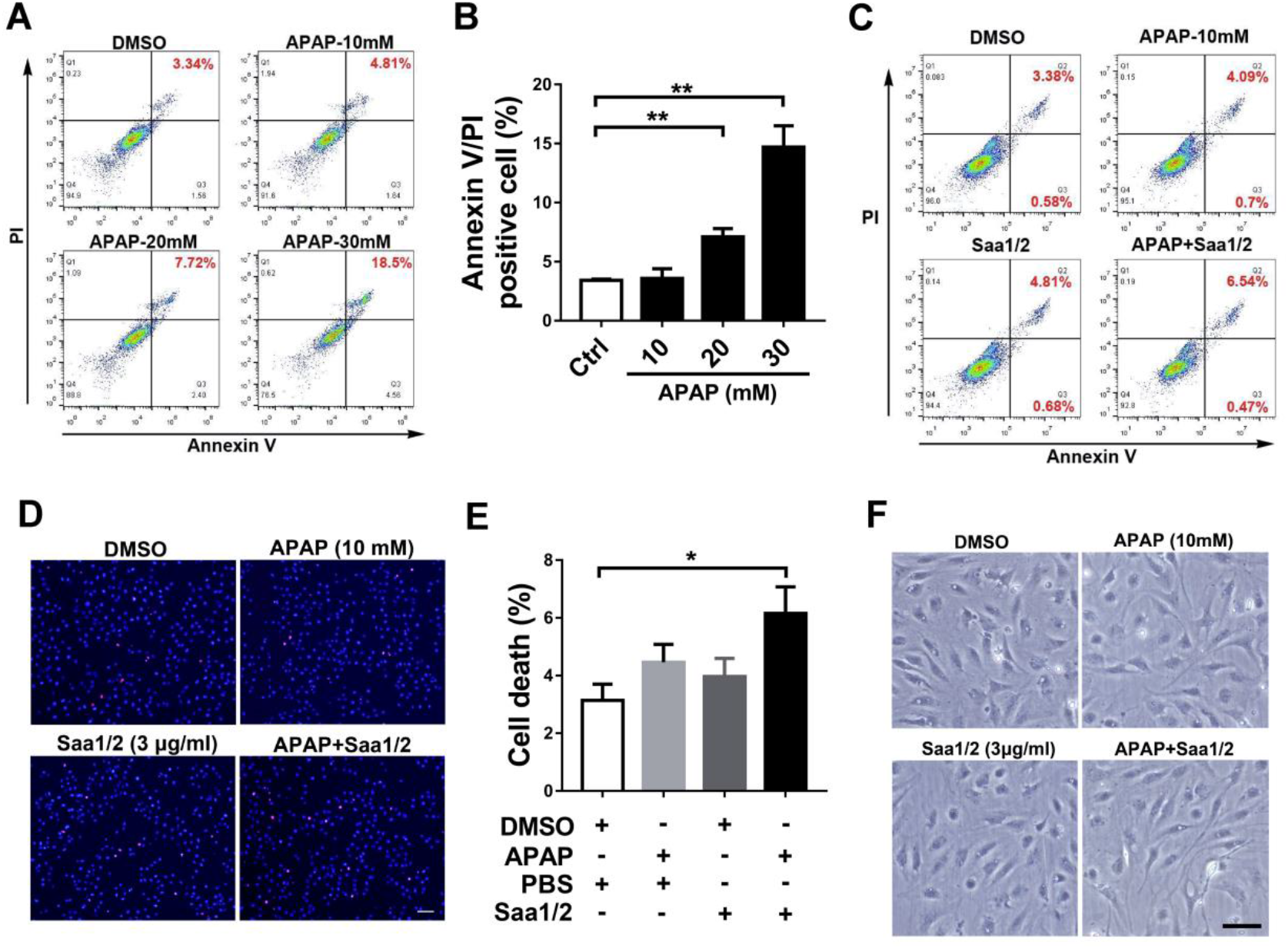
Saa1/2 promoted the injury effect of APAP on LSEC. LSECs were isolated from normal mice. (A) Flow cytometry was used to determine the proportion of dead LSECs induced by different concentrations of APAP (10, 20 or 30 mM) for 6 hours; (B) Quantification data of flow cytometry assay in (A) experiment (n=3); (C) Flow cytometry was used to determine the proportion of LSEC death induced by APAP (10 mM) or recombinant Saa1/2 proteins alone or mixed together; (D) Hoechst 33342 and PI double staining was used to measure the proportion of LSEC deaths induced by stimulating of APAP (10 mM) or recombinant Saa1/2 alone or combined together for 6 hours (scale bars = 50 μm); (E) Quantification data of double staining assay in (D) (n=3); (F) Cell morphology images of LSEC treated with APAP (10 mM) or Saa1/2 proteins alone or mixed together for 24 hours (scale bars = 50 μm). *, *p* < 0.05; **, *p* < 0.01.

### Saa1/2 promoted platelets adhesion to LSECs through Tlr2 receptor

In order to investigate the underling mechanism of Saa1/2 in promoting platelet aggregation in liver after APAP treatment, we firstly determined the expression of various Saa1/2 receptors and platelet adhesion-associated makers in LSECs under Saa1/2 stimulation (Supplementary Figure S3). Our data showed that Tlr2 and Vcam-1 were significantly induced by Saa1/2 in LSECs (Figure 5C and Supplementary Figure S3A). Furthermore, platelet adhesion experiments showed that Saa1/2 could significantly increase platelet adhesion to LSECs *in vitro*, while Cu-CPT22 (the TLR1/2 inhibitor) inhibited the effect of Saa1/2 on platelet adhesion (Figure 5A and B). Meanwhile, we also found that the induction of Vcam-1 expression by Saa1/2 was attenuated by Cu-CPT22 in LSECs (Figure 5C and D). And neutralizing Saa1/2 decreased the expression of Vcam-1 in liver tissues upon APAP treatment for 6 hours *in vivo* (Figure 5E and F). Finally, we treated LSECs with recombinant Saa1/2 alone or together with anti-Vcam-1 antibody, and found that blocking of Vcam-1 significantly reduced the adhesion of platelet on LSECs induced by Saa1/2 (Figure 5G and H). These findings suggest that Saa1/2 may promote platelets adhesion to LSECs through the Tlr2/Vcam-1 axis.

**Figure 5.**
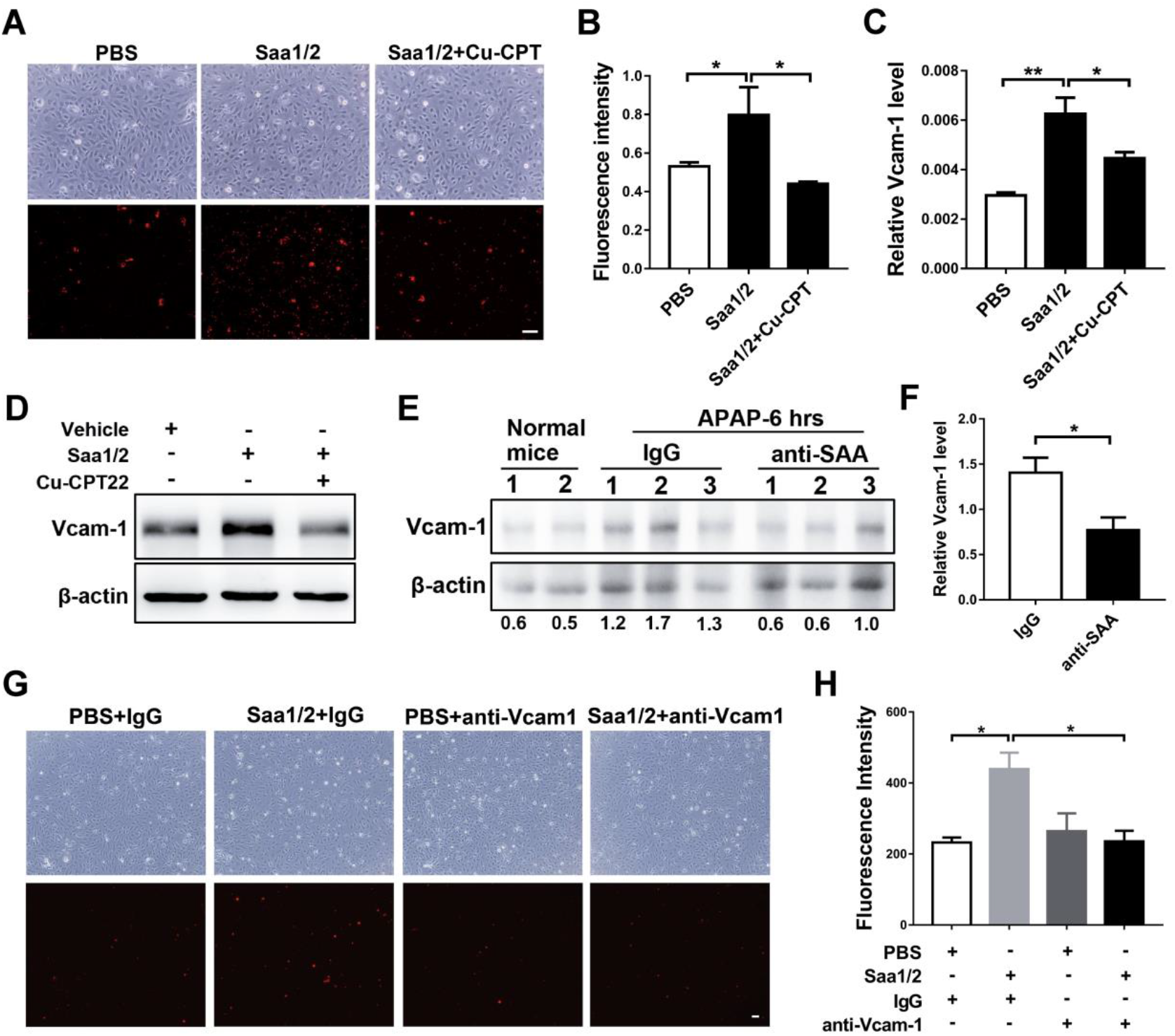
Saa1/2 promoted platelets adhesion to LSECs. (A) Representative images of adhesive platelets (red color). LSECs were cultured for 48 h in vitro and treated with Saa1/2 (3 ug/mL) alone or mixed with Cu-CPT22 (1 μM) for 2 hours, then Dil-labeled platelets were added and incubated with LSEC for 1 hour (scale bars = 50 μm); (B) Quantitative results of adhesive platelets in (A) (n=3); (C) Q-PCR analysis of Vcam-1 mRNA level in LSEC (n=3); (D) Western blotting analysis of Vcam-1 in LSEC; (E) Western blotting analysis of Vcam-1 in liver tissues, that were collected from mice treated with IgG or anti-SAA neutralizing antibody before injection of APAP (300 mg/kg) for 6 hours; (F) Quantification of relative Vcam-1 expression in (E) (n=3); (G) Representative images of adhesive platelets (red color) to LSECs treated with recombinant Saa1/2 alone or together with anti-Vcam-1 neutralizing antibodies (scale bars = 50 μm); (H) Quantitative results of adhesive platelets in (G) (n=3). *, *p* < 0.05; **, *p* < 0.01.

## Discussion

SAA is usually upregulated in inflammatory diseases and it could be as a sensitive marker as C-reactive protein (CRP) for diagnosis, prognosis and monitoring of tissue injury and infections [10]. The serum levels of SAA could be increased from 10- to 100-fold during limited inflammatory events to a 1000-fold increase during severe infections or inflammatory diseases [10, 17]. Recently, it has been reported that serum acute phase reactants, including SAA, were significantly upregulated in healthy individuals who are susceptible to APAP-induced liver injury [14]. In addition, serum SAA could also serve as a sensitive biomarker for early detection of hepatotoxicity induced by Ritodrine [15]. These findings suggested that SAA might involve in the pathogenesis of drug-induced liver injury. Therefore, monitoring the serum levels of SAA and other acute phase proteins prior to APAP treatment may help to avoid and manage the occurrence of drug-induced acute liver injury.

SAA is mainly produced and secreted by hepatocyte after tissue injury or infections. However, the expression and function of SAA in APAP and other drug-induced acute liver injury remain unclear. Although one study had declared that serum SAA was not statistically induced by APAP [15]. Here, we performed immunohistochemistry, real-time quantitative PCR, western blot and ELISA experiments to determine the expression of Saa1/2 in APAP-induced mouse liver injury model, our study showed that in the early stage (6-24 h) of acute liver injury induced by APAP in mice, the levels of Saa1/2 in liver tissue and serum were increased dramatically, and followed by gradually recovering to normal level at 24-72h after APAP injection (Figure 1A-D). Blocking the function of Saa1/2 either by neutralizing antibodies (anti-SAA) or small peptides (SAA-pep) significantly decreased the liver injury and necrosis areas, and more importantly increased the survival time of mice under the lethal dose injection of APAP (Figure 2). Moreover, our *in vitro* experiments also showed that the expression and secretion of Saa1/2 in primary hepatocytes were significantly upregulated under APAP stimulation. Therefore, based on our data, we believe that Saa1/2 can be induced by APAP and it may play an important role in APAP-induced acute liver injury.

Hepatic microvascular injury is an important early event in the pathogenesis of APAP-induced liver injury, and hepatic sinusoids injury preceded hepatotoxicity. As early as 30 minutes after the administration of APAP, the sinusoidal endothelial cells became swollen and large gaps formed by the destruction and coalescence of fenestrae within 2 hours [5, 7]. Then erythrocytes infiltrated into the space of disse via enlarged pores in sinusoidal lining cells at 2 hours, and the area occupied by red blood cells was markedly increased at 6 hours after APAP administration [6]. However, the mechanism of APAP-induced sinusoidal endothelial cell injury is still unclear. It has recently been reported that LSECs expressed CYP2E1, which is a key enzyme for metabolizing APAP into toxic intermediate NAQPI [18]. It suggests that APAP may directly induce sinusoid endothelial cell damage. In addition, TRAIL (tumor necrosis factor-related apoptosis-inducing ligand) can promote APAP-induced apoptosis of LSECs [19, 20]. Platelet adhered to hepatic sinusoidal endothelium promotes apoptosis of LSECs [21]. And kupffer cell is important for maintaining the integrity of hepatic sinusoidal endothelial cells [22, 23]. Moreover, it has also been found that inhibition of MMP-2 and MMP-9 activity can reduce APAP-induced sinusoidal endothelial cell injury [24]. Here in our study, we found that inhibition of Saa1/2 significantly reduced intrahepatic hemorrhage (Figure 3A-B), maintained the integrity of hepatic sinusoid structure (Figure 3C-E), and decreased the platelet accumulation in liver tissues from APAP-induced injury mice (Figure 3F-G). Our *in vitro* experiments data showed that Saa1/2 could promote APAP-induced LSECs cell death (Figure 4), but we did not observe the induction of Cyp2e1 by Saa1/2 in LSECs (Supplementary Figure S4), which indicated that Saa1/2 did not influence the metabolism of APAP in LSECs. Taken together, our findings suggest that, on the one hand, Saa1/2 may cooperate with APAP to directly promote LSECs injury, and on the other hand, Saa1/2 may induce cell death of sinusoidal endothelial cells by promoting platelet aggregation in liver. However, the exact mechanism of APAP induced hepatic sinusoidal endothelial cell injury is further needed to be clarified.

Intrahepatic platelet agglutination is an obvious feature during the process of APAP-induced liver injury. Some studies have shown that inhibition of platelet activation by Lepirudin (a thrombin inhibitor) or depletion of platelet by anti-CD41 neutralizing antibodies could significantly reduce the severity of APAP-induced liver injury [25]. In addition, Chi3l1 (Chitinase 3-like-1) promoted liver injury by recruiting platelets into liver in APAP-induced liver injury model. Accordingly, specific monoclonal antibody against Chi3l1 could effectively inhibit intrahepatic platelet accumulation and alleviate liver injury caused by APAP [26]. However, little is known about the molecular mechanism by which platelets were recruited to the liver during APAP-induced acute liver injury. Our study found that inhibition of Saa1/2 by neutralizing antibodies significantly reduced the deposition of fibrin(ogen) in liver tissue (Supplementary Figure S5). As fibrin is a key factor in mediating platelet recruitment and aggregation, we speculate that Saa1/2 may inhibit platelet recruitment by reducing the deposition of fibrin in liver. In addition, our *in vitro* experiments showed that Saa1/2 promoted Vcam-1 expression in LSECs and induced platelet adhesion to LSECs through Tlr2/Vcam-1 axis (Figure 5A-B).

In conclusion, our findings cleared that Saa1/2 expression was up-regulated in the early stage of APAP-induced acute liver injury (within 24 h), and inhibition of Saa1/2 could significantly reduce intrahepatic bleeding, maintain the integrity of hepatic sinusoids and decrease platelet accumulation in the liver, thus lead to mitigation of liver injury. Therefore, SAA may be a potential therapeutic target for APAP-induced acute liver injury.

## Materials and Methods

### Animals and APAP model

BALB/c mice were purchased from Beijing Vital River Laboratory Animal Technology Co., Ltd, China. Mice were housed under a 12 h light-12 h dark cycle, allowed unrestricted access to standard mouse chow and tap water, and acclimated to these conditions for at least 1 week before inclusion in experiment. This study was in accordance with the animal welfare and ethics requirements of The Animal Ethics Committee of Guangzhou Institute of Biomedicine and Health, Chinese Academy of Sciences. Acute liver injury was induced by APAP in 6-8 weeks old BALB/c mice, which were fasted for 16 hours before a single intraperitoneal injection with 300 or 500 mg/kg APAP. Subsequently, mice were sacrificed at different time points (6 hours, 1 day, 3 days and 7 days). In the Saa1/2 neutralizing experiment, the neutralizing antibody (anti-SAA) or small peptide (SAA-pep) were intravenously injected in mice one hour before APAP injection. The anti-SAA antibody was purchased from Abcam (cat. ab124964). The SAA-pep (YIGSDKYFHARGNY), which sequence was identical with human SAA1/2 at aa29-42, and the Scramble (NYAGRKFHYSGDYI) were synthesized by Sangon Biotech Co., Ltd, Shanghai, China.

### Isolation of mouse primary hepatocytes (MPH) and liver sinusoidal endothelial cells (LSEC)

MPH and LSEC were isolated from male mice at aged 8-10 weeks by collagenase perfusion and density gradient centrifugation. Briefly, mice were firstly anesthetized by 3-bromoethanol (TBE) solution according to their body weight (400 mg/kg). The liver was perfused with collagenase solution (0.1 mg/mL) for 20 minutes. The digested liver was removed, minced and filtered through 100-μm and 40-μm strainers in turn to obtain single cell suspension. For MPH isolation, the cell suspension was centrifuged at 50 × *g* and washed with DMEM for 3 times, and the cell precipitates were re-suspended with medium (DMEM+10%FBS+1%ITS+0.1μM dexamethasone+ 1% penicillin and streptomycin) and cultured in collagen-coated plates at 37°C and 5% CO_2_ conditions. For LSEC isolation, after repeated centrifugation at 50 × *g*, the supernatant that containing the non-parenchymal cell were centrifuged at 350 × *g* for 10 min. Cell pellets were re-suspended in 16% (Iodixanol, w/v) OptiPrep solution (cat: AXS-1114542, Serumwerk Bernburg, Germany) and top with 11.5% OptiPrep and then with 1 × PBS. After centrifugation at 850 × *g* for 20 min, the visible cells between layers of 16% and 11.5% OptiPrep were collected. Cells were washed and re-suspended with warm RPMI1640+10% FBS, and the LSECs were separated from KCs by placing cells in a polystyrene Petri dish and incubated in a humidified culture incubator for 8 min at room temperature. LSECs were obtained and cultured in RPMI1640+5% FBS+1% penicillin and streptomycin at 37 °C and 5% CO_2_ conditions.

### Analysis of gene expression

Total RNA was extracted from liver tissues or cultured cells by Trizol reagent (Med Chem Express). The relative mRNA expression of genes was analyzed by real-time quantitative PCR (q-PCR) with Bio-Rad CFX96. Gapdh was used as an internal control. Sequences of all primers were listed in Supplementary Table 1.

Western blotting was performed to detect the protein level of Saa1/2 (cat. ab124964; Abcam), β-actin (cat.4970; Cell Signaling Technology) and Vcam-1 (cat.32653S; Cell Signaling Technology).

### Enzyme-linked immunosorbent assay (ELISA)

The serum levels of Saa1/2 were determined by mouse SAA ELISA kit (cat.EK1190, Boster Biological Technology co.Itd, China). The procedure was carried out according to the manufacturer’s instructions. The mouse serum samples were obtained from heart and detected immediately or frozen at −80°C.

### Histology and immunohistochemistry

Mouse liver tissues were fixed in 4% paraformaldehyde (PFA) for 24 hours, embedded in paraffin and cut into 5-μm-thick sections and then stained with hematoxylin and eosin (H&E). For immunohistochemistry staining, antigen retrieval was performed by incubating slices with citrate-EDTA buffer (cat.P0086, Beyotime, China) and heated at high fire in a microwave oven for 5 min, followed by 30 min of cooling at room temperature. Then endogenous peroxidases were quenched by incubating the slices with 0.3% H_2_O_2_ for 10 min, and slices were blocked with10% FBS for 1 hour followed by incubation with anti-SAA (cat.ab124964, Abcam) or anti-Cd42b (cat.ab183345, Abcam) primary antibody at 4°C overnight. Secondary antibody conjugated with horseradish peroxidase (HRP) was incubated at 37°C for 30 min. Then Color development was achieved by DAB substrate (cat.K3468, DAKO). Sections were then counterstained with hematoxylin, dehydrated through ethanol and xylene, and mounted by using a xylene-based mounting medium.

### Immunofluorescence

Mouse liver tissues were fixed in 4% PFA for 24 hours, and cut into 20-μm-thick sections by Vibratomes (Leica Biosystems), and then slices were blocked with10% FBS and 0.1% Triton X-100 for 2 hours. Subsequently, slices were incubated with anti-hepatic sinusoidal endothelial cell specific marker SE-1 primary antibody (cat.NB110-68095; NOVUS, 1:100) overnight at 4°C. Rinsed 3 times with 1 × PBS, and incubated with secondary antibody for 1 hour at room temperature. Nuclei were stained with DAPI for 5 min. Finally, the images were observed using laser scanning confocal microscope (Zeiss 710 NLO, Germany).

### Evans Blue assay

The permeability of hepatic sinusoidal endothelium was examined by Evans Blue. Evans blue dye was dissolved into normal saline (5 mg/mL) and injected into mice via tail vein at a concentration of 30 mg/kg one hour before sacrificed. The liver tissue was divided into two parts, one part of liver were fixed in 4% PFA and cut into 20-μm-thick sections for fluorescence analysis of Evans Blue signaling by using confocal microscope. And another part of liver tissue was weighted, minced and dissolved in formamide (1 g/ml) and incubated in a water bath at 55°C for 24 hours. Then the samples were subjected to centrifugation at 4°C, 5000 × *g* for 30 min, and the supernatants were transferred to 96-well plate for measuring Evans Blue concentration with a spectrophotometer at 620/680nm.

### Analysis of cell death

In this study, Annexin-V/PI apoptosis detection kit (cat.556547, Becton Dickinson) and Hoechst33342/PI double staining kit (cat.KGA212, KeyGEN BioTECH, China) were used to evaluate the cell death rate of LSEC that were treated by APAP or Saa1/2. The procedure was carried out according to the manufacturer’s instructions.

### Platelet adhesion assay

Firstly, the platelets were separated with steps as follow. In brief, blood sample (1 mL) was collected from heart using an anticoagulant syringe containing 150 μL ACD buffer, added 3 mL washing buffer into sample and centrifuged at 250 × *g* for 10 min. The supernatant was transferred to a new 15 mL polypropylene conical tube to remove the red and white blood cell pellet, then added washing buffer to 10 mL and 0.1 μM PGE1 to inhibit platelet activation, centrifuged at 1250 × *g* at room temperature for 10 min. The platelet pellet was washed by washing buffer containing 0.1 μM PGE1 twice. Then platelets were incubated with Dil stain (5 μM) for 10 min at room temperature, centrifuged at 1250 × *g* for 10 min and added to LSECs on the culture plates. After incubation with LSEC for 1 hour, the unattached platelets were washed by PBS, and LSECs were fixed by 4% PFA for photographing. The fluorescence intensity of Dil-labeled platelets was analyzed by Image J software (version 1.52a). The reagents were prepared as follows: 1) ACD buffer (pH=4.5): 85 mmol/L trisodium citrate, 71 mmol/L citric acid and 111 mmol/L dextrose; 2) washing buffer: 5.36 mmol/L NaCl, 0.12 mmol/L KCl, 0.012 mmol/L NaH_2_PO_4_.H_2_O, 0.2 mmol/L HEPES, 0.2 mmol/L glucose, 0.08 mmol/L MgCl_2_, 33.33 mmol/L EGTA, 5% NaHCO_3_ and 0.08% BSA.

### Statistical analysis

Each experiment had at least three independent experiments, and the data were expressed by mean ± SEM. Data of two groups was analyzed by unpaired student’s-*t* test, data of more than two groups was analyzed by one-way or two-way ANOVA followed by Dunnett’s or Sidak’s test, respectively. Survival curve was compared using log-rank (Mantel–Cox) test. All statistical analyses were performed using GraphPad Prism software (version 7.0). All statistical tests were two-sided, and p < 0.05 was considered to be statistically significant.

## Acknowledgements

The authors wish to acknowledge outstanding technical support from the animal center and instrument center of Guangzhou Institutes of Biomedicine and Health, Chinese Academy of Sciences. This work was supported by the National Key R&D Program of China (2019YFA0111300), Guangzhou City Science and Technology Plan (202102021138), the National Natural Science Foundation of China (31871379, 31900539), Guangdong Province Science and Technology Plan (2018A050506070), the Sino-German rapid response funding call for COVID-19 related research (C-0031) and Frontier Research Program of Bioland Laboratory (Guangzhou Regenerative Medicine and Health Guangdong Laboratory) (2018GZR110105011).

