## Supplementary Figures and Table for "Serum Amyloid A promotes Acetaminophen-induced liver injury by damaging sinusoidal endothelial cell and exacerbating platelet aggregation in liver"

### Supplemental Figures

*Figure S1*

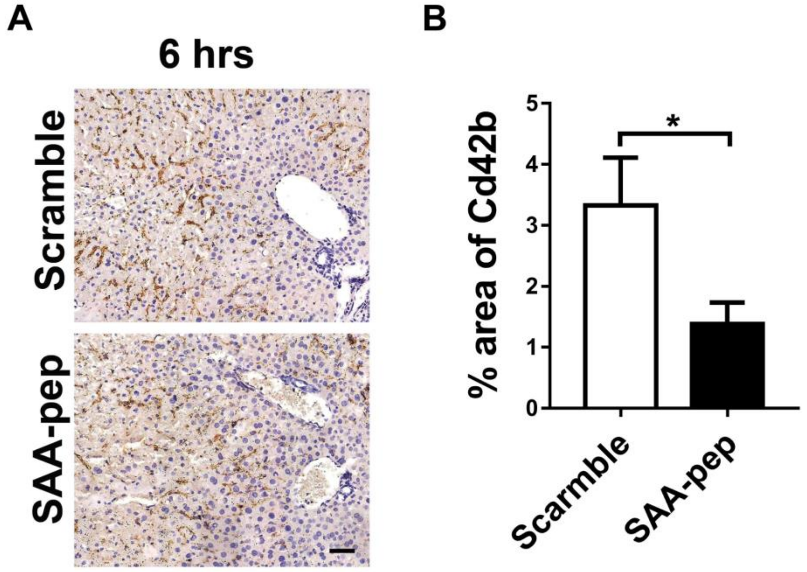


**Figure S1. Treatment of SAA-pep decreased intrahepatic platelet accumulation.** Mice were treated with neutralizing peptide derived from human SAA1 (aa29-42) or the scramble peptide (50 μg/mice) one hour before APAP-induced mouse liver injury. (A) Immunohistochemistry staining of platelet with Cd42b antibody (scale bar = 50 μm), (B) and quantitative data of Cd42b positive areas in (A). *, $p < 0.05$.

*Figure S2*

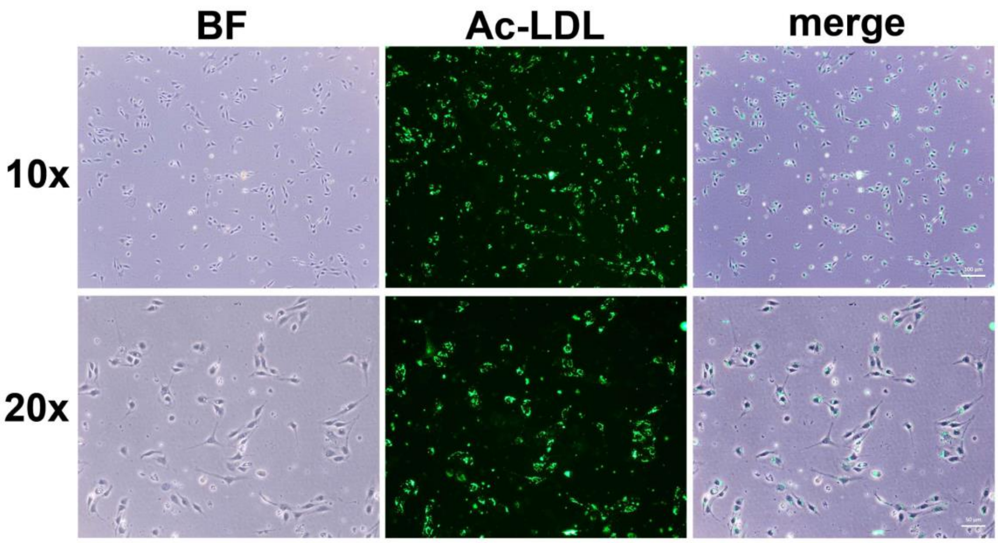


**Figure S2. Identification of primary liver sinusoidal endothelial cells.** liver sinusoidal endothelial cells (LSECs) were isolated from normal mice, and cultured in RPMI-1640 medium with 5% fetal bovine serum for 24 hours. The purity of isolated LSCEs was more than 94%, while assessed by up taking acetylated low-density lipoproteins (Ac-LDL).

*Figure S3*

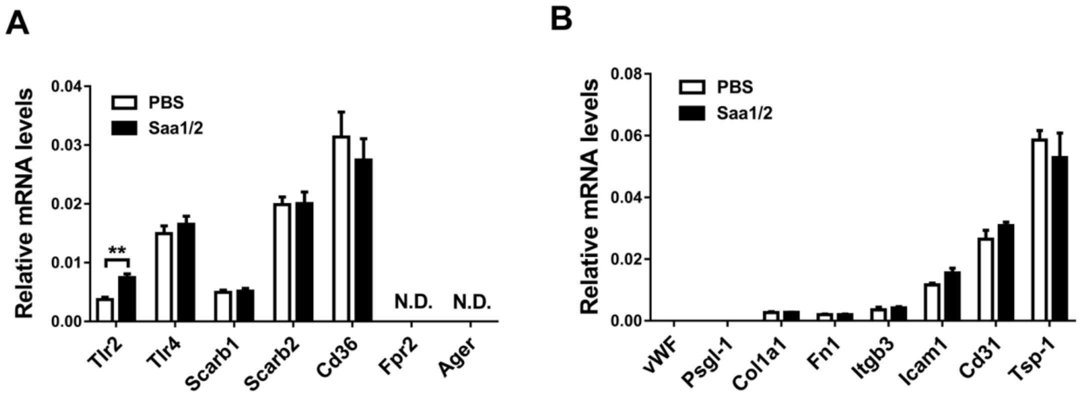


**Figure S3. Saa1/2 induced Tlr2 expression in LSECs.** LSECs were treated recombinant Saa1/2 proteins (3 μg/mL) for 2 hours, then the expression of Saa1/2 receptors (A) and platelet adhesion-associated genes (B) was detected by q-PCR. N.D. means not detected by q-PCR.

*Figure S4*

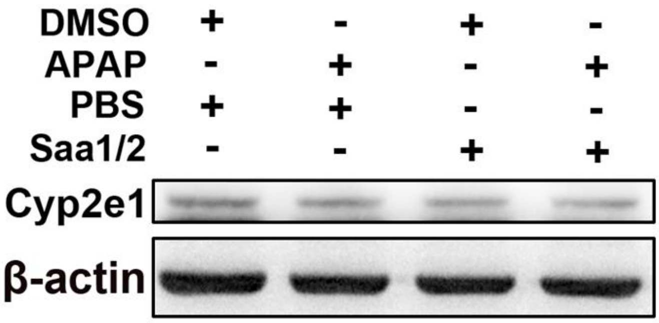


**Figure S4. Saa1/2 and APAP did not induce Cyp2e1 expression in LSECs.** LSECs were treated with APAP (10 mM) or recombinant Saa1/2 proteins alone or mixed together for 6 hours, and the expression of Cyp2e1 was determined by western blotting.

*Figure S5*

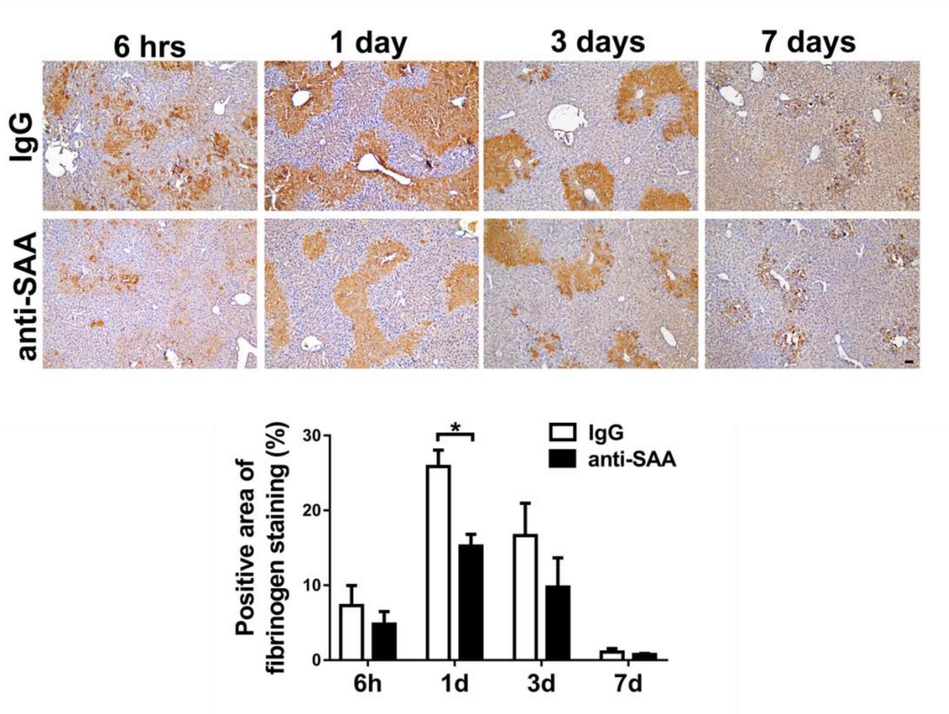


**Figure S5. Treatment of anti-SAA decreased intrahepatic fibrinogen deposition.** Mice were treated with Saa1/2 neutralizing antibodies (3 μg/mice) one hour before APAP-induced mouse liver injury, and scarified at the indicated time points. Images of immunohistochemistry staining of fibrinogen (scale bar = 50 μm) and quantitative data of fibrinogen positive staining areas were shown. *, $p < 0.05$.

### Supplemental Table

**Table S1. Primers for qPCR analysis**

| Gene name | Sense strand (5'-3') | Antisense strand (5'-3') |
|---|---|---|
| Saa1/2 | GCGAGCCTACACTGACATGA | TGCCGAAGAATTCCTGAAAG |
| Tlr2 | CTCTTCAGCAAACGCTGTTCT | GGCGTCTCCCTCTATTGTATTG |
| Tlr4 | ATGGCATGGCTTACACCACC | GAGGCCAATTTTGTCTCCACA |
| Scarb1 | CCCCAGGTTCTTCACTACGC | TCCTTATCCTGGGAGCCCTT |
| Scarb2 | TCTTCCCGTCAGACTTGTGC | AGTCCATGCAGTTTCCCTCG |
| Cd36 | ATGGGCTGTGATCGGAACTG | GTCTTCCCAATAAGCATGTCTCC |
| Fpr2 | TTACACCACAGGAACCGAAGA | AAGTGATGGAGACAACCACCA |
| Ager | CTTGCTCTATGGGGAGCTGTA | CATCGACAATTCCAGTGGCTG |
| Vwf | GATTGCCTGTGTAACGCAGTA | GCAGGTCAGGTTGCAGGAAT |
| Psgl-1 | GGGATGGTCCTTCCTTTGGG | ACAATGGTCTAAGCGCCCTC |
| Col1a1 | ATGATGCTAACGTGGTTCGT | TGGTTAGGGTCGATCCAGTA |
| Fn1 | ATGAGCGCCCTAAAGATTCC | CCAGCAGCATGATCAAAACA |
| Itgb3 | GGACAACTCTGGGCCGCTC | GGTTACATCGGGGTGAGCC |
| Icam-1 | CTGGGCTTGGAGACTCAGTG | CCACACTCTCCGGAAACGAA |
| Cd31 | GACCCCCAGAACATGGATGTAG | TAAGGGAGCCTTCCGTTCTTAG |
| Tsp-1 | GGAGGAGCTGTTCGGTACTA | AAAGGGAGAAAGTCCAGAAGCTAT |
| Vcam-1 | CTCTTCAGCAAACGCTGTTCT | GGCGTCTCCCTCTATTGTATTG |
| Gapdh | AACTTTGGCATTGTGGAAGG | CACATTGGGGGTAGGAACAC |
